# OPM-FLUX: A Pipeline for OPM MEG Data Analysis

**DOI:** 10.64898/2026.04.24.720604

**Authors:** Arnab Rakshit, Tara Ghafari, Anna Kowalczyk, Ole Jensen

## Abstract

Optically pumped magnetometer-based magnetoencephalography (OPM-MEG) has recently emerged as a powerful neuroimaging approach in cognitive neuroscience, extending beyond the limitations of conventional cryogenic systems with greater experimental flexibility and wearable recording. Despite these advantages, standardised data analysis frameworks specifically tailored to OPM technology are still lacking, leading to variability in processing choices and reduced reproducibility across laboratories and hardware platforms. We introduce OPM-FLUX, a comprehensive and fully documented end-to-end analysis pipeline developed for OPM-MEG data. The pipeline defines a clear sequence of preprocessing, noise suppression, artifact handling, spectral analysis, evoked response analysis along with recommended parameter settings. It also includes source reconstruction to identify where in the brain the signals originate. In addition, OPM-FLUX supports multivariate pattern analysis (MVPA), enabling time-resolved decoding of cognitive processes from sensor level data. OPM-FLUX is implemented in MNE-Python and distributed as interactive Jupyter Notebooks that combine executable code with detailed methodological explanations and graphical outputs. The pipeline further provides standardized reporting templates and a data acquisition Standard Operating Procedure to facilitate preregistration, consistent documentation, and standard practices across research sites. The workflow is demonstrated using openly available datasets acquired from both Cerca/QuSpin and FieldLine OPM systems during a visuospatial attention paradigm that modulates alpha, beta, and gamma oscillations and elicits event-related responses. By supporting multiple OPM platforms and promoting consistent methodological choices, OPM-FLUX enhances transparency, comparability, and replication in OPM-MEG research. The pipeline also serves as an educational resource for students and researchers entering the field and is designed to evolve alongside ongoing technological and methodological advances in OPM-based brain imaging.

## 1. Introduction

Optically pumped magnetometer-based magnetoencephalography (OPM-MEG) has recently emerged as a powerful brain imaging modality in cognitive neuroscience, as demonstrated across multiple OPM-MEG studies (Boto et al., 2018; Sander et al., 2012; Schwindt & Johnson, 2010; Xia et al., 2006) extending the capabilities of traditional MEG (Baillet, 2017) by enabling high spatial and temporal resolution with greater experimental flexibility (Brookes et al., 2022; Hill et al., 2020). MEG research relies on extensive signal processing to interpret the magnetic fields generated by neuronal activity, and the field has greatly benefited from the development of numerous open-source analysis toolboxes and processing pipelines (Baillet, 2017; Ferrante et al., 2022; Gramfort et al., 2013; Hinkley et al., 2020; Litvak et al., 2011; Oostenveld et al., 2011; Tadel et al., 2011). which have great potential in translational neuroscience research (Brickwedde et al., 2024) and offer several advantages over traditional cryogenic MEG including improved signal quality due to the closer proximity of sensors to the scalp, enhanced participant mobility, and reduced operational costs that facilitate wider adoption and more naturalistic experimental paradigms. Hence there is a growing need for dedicated signal processing and analysis pipelines tailored specifically to the unique characteristics of OPM-based measurements.

Since OPM is a relatively new technology, a clear ‘best-practice framework’ has not yet been established. Therefore, it is crucial to converge on the fundamental signal processing steps of OPM-MEG data that should be followed and to standardise them. The existing pipelines, originally developed for conventional MEG data must be adapted to OPM-MEG data analysis as there are differences in sensor technology and operating principles. Although a few toolboxes have attempted to offer basic functionalities, such as noise suppression and artifact rejection, co-registration with MRI (Cao et al., 2022; Takeda et al., 2019), some depend on licensed proprietary software environments, which may limit accessibility, reproducibility, and community-driven development. There remains a need for a fully open-source standard for OPM data processing that supports the complete analysis pipeline. Moreover, multiple methods exist for performing the same step, each requiring several parameter adjustments. This lack of uniformity creates challenges for comparing and replicating results across studies and systems.

We here introduce an end-to-end signal processing pipeline, named OPM-FLUX, specifically designed for OPM-MEG. Here, “end-to-end” refers to a complete workflow that covers the main stages of OPM-MEG data handling, from data acquisition and preprocessing through to analysis and reporting. The pipeline provides the essential processing steps required for conducting OPM-based cognitive neuroscience experiments, along with detailed explanations for each step, while adhering to the general guidelines established for MEG research (Hari et al., 2018). The pipeline is implemented using the MNE-Python toolbox (Gramfort et al., 2013)although the procedures and recommendations can be adapted to other toolboxes. OPM-FLUX is developed and validated using data acquired from both Cerca/QuSpin and FieldLine OPM systems; however, the intent is to extend it to other OPM platforms in the future. By supporting multiple hardware implementations, the pipeline will facilitate replication and comparison of results across research sites employing different OPM-MEG systems. The use of an open-source framework ensures that researchers worldwide can access and utilize the pipeline thereby promoting inclusivity, transparency, and reproducibility in OPM-MEG research.

The replicability of any research depends on clearly describing the processing and analysis steps used to reach the conclusions, making it essential to detail these procedures in the manuscript.

Currently, researchers around the world often use different approaches and parameter settings for performing similar analysis step, which makes reproducibility challenging and leads to considerable variation in the methods sections of published studies. To address these issues, OPM-FLUX provides a set of specific standardized processing steps with optimized parameter settings, allowing researchers to follow a common and consistent pipeline. The pipeline also provides specific recommendations and text for how to report the parameters and algorithms used in the analysis. In summary, the standardized approach we here propose facilitates a collaborative and coherent research ecosystem, where future studies can reliably build upon the results and methodologies established by previous work.

In the context of modern open-science practices, where pre-registration is becoming increasingly common (Shrout & Rodgers, 2018), the proposed OPM-FLUX pipeline offers significant advantages by simplifying the process of outlining detailed analysis steps prior to conducting a study. The pipeline includes ready-to-use text suggestions that can be directly incorporated into the methods section of a manuscript or a pre-registration document, thereby saving time and ensuring consistency. Additionally, a Standard Operating Procedure (SOP) for the OPM data acquisition process is provided with the pipeline, promoting standardized data acquisition practices and supporting uniform operational protocols across laboratories. The OPM-FLUX pipeline is well aligned with current best practices and methodological frameworks presented in recent OPM publications (Boto et al., 2018, 2019; Brickwedde et al., 2024; Brookes et al., 2022). It adheres to the established guidelines for MEG research reproducibility, helping to ensure transparent, reliable, and replicable scientific outcomes (Gross et al., 2013; Hari et al., 2018; Pernet et al., 2020).

Another key objective of the proposed OPM-FLUX pipeline is to support education and training in OPM-based cognitive neuroscience. The OPM technology offers several advantages over traditional cryogenic MEG, including improved signal amplitude due to the closer proximity of sensors to the scalp, greater mobility, and lower recurring costs. These benefits are gradually shifting the focus of cognitive neuroscience research toward OPM systems, leading to a growing number of researchers worldwide adopting this technology. The OPM-FLUX pipeline, with its comprehensive descriptions of processing steps and parameter settings, serves as an excellent educational resource for students and new researchers entering the field.

In summary, OPM-FLUX aims to provide a standardized, transparent, and accessible framework for OPM-MEG data acquisition, processing, reporting, and education. By promoting consistency across laboratories and hardware systems, the pipeline is intended to improve reproducibility, support open-science practices, and facilitate the broader adoption of OPM-MEG in cognitive and translational neuroscience.

## 2. Approach

### 2.1 Dataset

There is a growing need to share recorded OPM datasets with the broader scientific community to enhance accessibility and transparency (Niso et al., 2016).To support this objective, OPM-FLUX offers two high-quality example datasets that are freely accessible via the Neural Oscillation Group website (https://www.neuosc.com/flux) recorded using a Cerca and a Fieldline system. At a later stage more datasets will be included to conduct group analysis.

### 2.2 Implementation

The OPM flux pipeline is developed as interactive scripts using Jupyter notebooks. This allows for demonstrating the code line-by-line in combination with detailed descriptions and graphical outputs. Ultimately users can copy the code and integrate it into standard Python scripts for batch processing. The pipeline uses MNE-Python toolbox, however custom functions are also introduced in some of the scripts. The scripts for the Cerca and Fieldline dataset are openly available linked via https://www.neuosc.com/flux.

Contrary to other popular toolboxes being based on Matlab e.g. FieldTrip (Oostenveld et al., 2011), Brainstorm (Tadel et al., 2011) and SPM (Litvak et al., 2011), the pipeline is based on MNE-Python for the following reasons: first, Python is accessible without license (in contrast to Matlab); second, MNE-Python is constantly improved by an active community and third, it has gained strong popularity among MEG researchers worldwide.

### 2.3 The hardware of the OPM system

To demonstrate the workflow of the pipeline, we use a dataset recorded using the Cerca/Quspin system at OPM facility at Oxford Centre for Human Brain Activity (OHBA), University of Oxford. The dataset was recorded from a participant performing a visuospatial attention task. In addition, OPM-FLUX has been tested on datasets acquired using a FieldLine OPM-MEG system at the Centre for Human Brain Health (CHBH), University of Birmingham.

The Cerca system uses optically pumped magnetometer (OPM) sensors from QuSpin (QZFM Gen-3) which are capable of measuring magnetic fields along three orthogonal axes (X, Y, and Z Fig. 1A). In contrast, the FieldLine HEDscan system equipped with V3 dual axis sensors measure magnetic fields along Z and Y directions; however, at the time of the experiment acquisition software allowed measurement of magnetic activity only along the Z-axis, which is oriented approximately perpendicular to the scalp (Fig. 1B).

**Figure 1.**
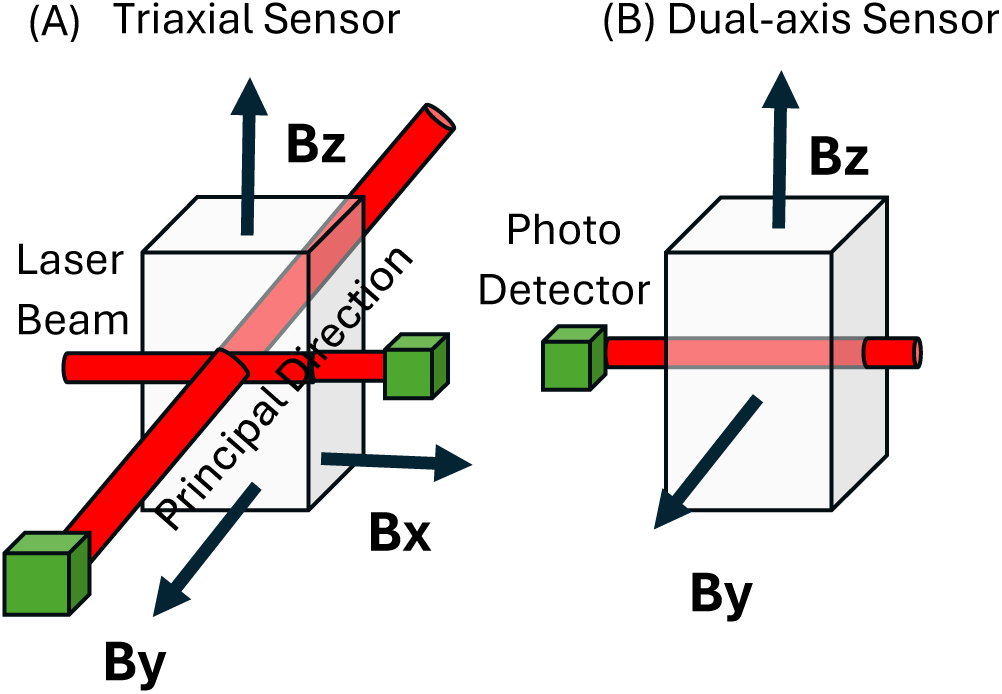
Triaxial and single-axis sensors used in the Cerca and FieldLine systems, respectively. (A) The triaxial sensor employs two laser beams and effectively enables measurement of magnetic field components along three orthogonal axes. (B) The dual-axis sensor employs a single laser beam and is sensitive to magnetic field components perpendicular to the beam direction (Bz and By).

In the triaxial QuSpin sensor design, a single 795 nm laser is split into two beams that pass through the same ^87^Rb vapour cell (Osborne et al., 2018). These two optical channels, in combination with applied orthogonal magnetic-field modulations and phase-sensitive demodulation, allow reconstruction of all three components of the magnetic field vector. Each optical channel is primarily sensitive to two magnetic-field components transverse to the beam direction rather than to the component parallel to the beam. Consequently, triaxial magnetic field reconstruction does not rely on the assumption that longitudinal magnetic fields produce no measurable effect, but instead emerges from the specific optical geometry modulation/demodulation scheme (Boto et al., 2022)

By comparison, the FieldLine sensor employs a single laser beam and detects magnetic fields perpendicular to the laser beam propagation direction, (corresponding to the Z axis in the present setup;(Alem et al., 2023)). As noted above, future software updates on the HEDscan system are expected to extend this capability to dual-axis sensing, with the third magnetic-field component estimated via interpolation.

Sensors from both OPM-MEG systems were operated in a closed-loop mode. In this mode, on-board compensation coils are dynamically driven by the demodulated photodiode signal to null the magnetic field at the vapour cell. The feedback current required to maintain the sensor near the zero-field resonance condition is used as the output signal.

### 2.4 The Paradigm

The paradigm requires the participant to detect the appearance of a small dot in the centre of one of the two moving gratings (Fig. 2). During the paradigm a fixation dot always remain on the screen and a cue pointing towards the left/right direction appears at the beginning of each trial. The cue indicates which of the two stimuli (moving gratings in left and right hemifields) should be attended. After a 1s inter-stimulus interval, the two moving gratings appear on the screen and after a random interval of 0.1 – 1.9s a white dot appears for 0.05s in the centre of the cued stimulus. The moving gratings remain on the screen for a total of 3 s. The participants are instructed to press a key using their right index finger when they detect the cued white dot. The button must be pressed within 1.5 s to be considered as a valid response. To ensure that participants are engaged, there are 10% catch trials in which the white dot does not appear. A total of 6 blocks of 60 trials were repeated. After ending of each block, the accuracy score is shown to the participants. The paradigm was presented using a python based open-source toolbox named PsychoPy (Peirce et al., 2019)

**Figure 2.**
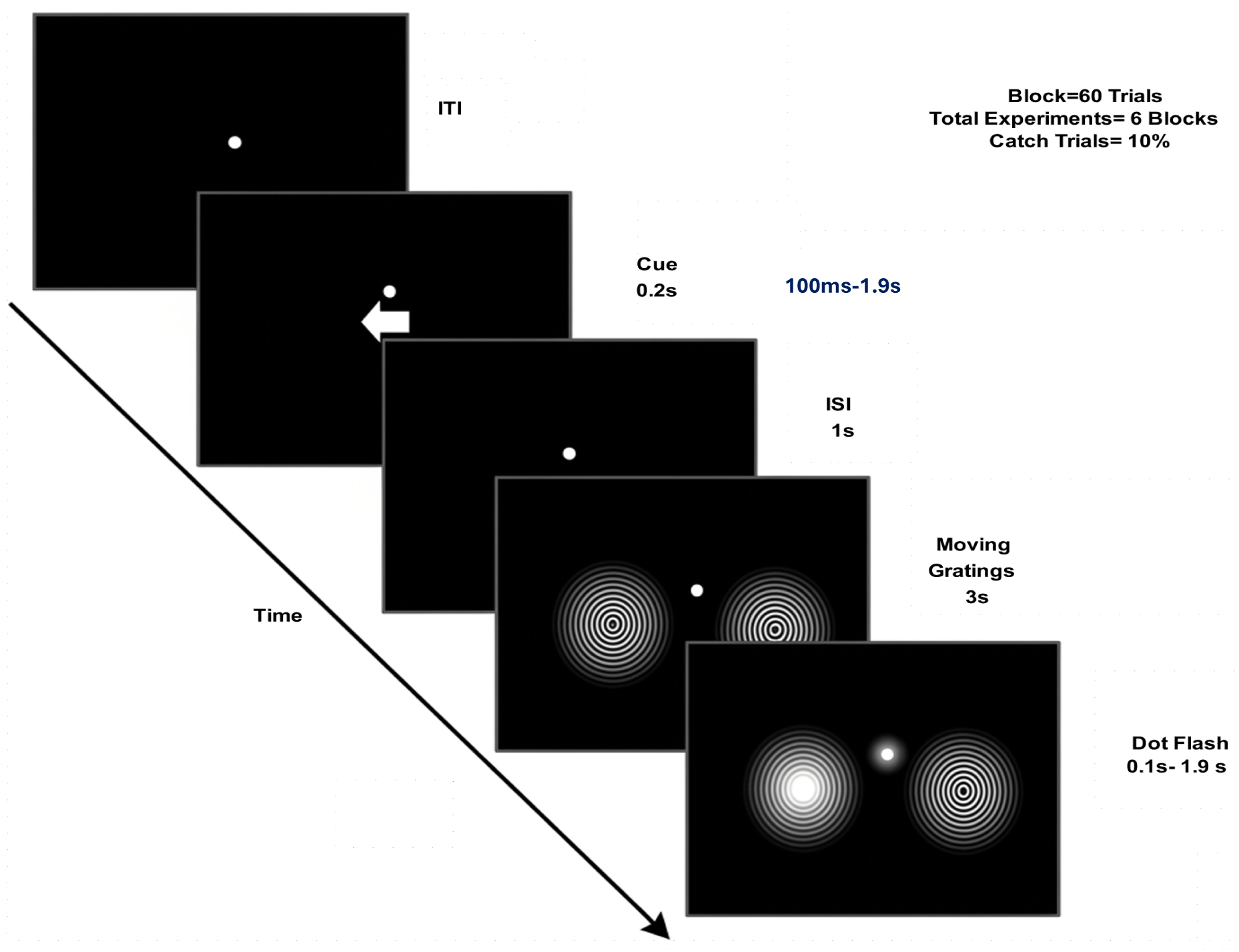
The experimental paradigm used for OPM/MEG data collection in the FLUX pipeline. Each trial begins with a cue instructing participants to attend to either the left or right visual field, producing clear hemispheric lateralisation of posterior alpha power (8 – 12 Hz). This is followed by the presentation of moving gratings, which elicit strong gamma-band activity (60 – 90 Hz) and visual evoked fields. Finally, a button press in response to a dot flash generates gamma activity in the motor cortex, followed by beta-band (15 – 30 Hz) modulation. Because this task reliably induces well-known modulations in alpha, beta, and gamma oscillations, as well as distinct event-related fields, it serves as a suitable test paradigm for validating and demonstrating the capabilities of the FLUX pipeline.

The paradigm is chosen as it typically modulates the alpha (8 -12 Hz) activity in posterior cortex according to the spatial cue (Jensen & Bonnefond, 2026) the grating will induce gamma (60 -90 Hz) activity in the visual cortex (Hoogenboom et al., 2006) and the motor response is expected to modulate the beta (15 -30Hz) activity in the motor cortex (Cheyne, 2013). Furthermore, even-related fields will be evoked by the cue and grating onsets.

## 3. Standard Operating Procedures

A standard operating procedure (SOP) for OPM data acquisition can be found on neuosc.com. It includes every necessary step for conducting an OPM experiment. Although the steps are specifically applicable for Cerca/Quspin system it generalises to other OPM systems as well. It is important to combine the SOP and analysis pipeline in a single framework to ensure appropriate channel naming and procedural consistency.

### 3.1 Co-registration

Co-registration was used to determine the position and orientation of the sensors relative to each participant’s head thereby enabling accurate source reconstruction.

In the Cerca system, head shape digitisation was carried out using an EinScan infrared 3D scanner (ref?), with texture flashing disabled to ensure participant comfort and safety. The participant was seated and asked to remain still during scanning. Digitisation was performed twice, once with the participant wearing the helmet immediately after data acquisition, and once after the helmet was removed. Anatomical fiducial landmarks (nasion, left and right Helix-Tragus Junction points; Fig. 3) and few facial points were subsequently identified and selected by the experimenter on the acquired 3D image using the 3D processing software. These fiducials were used to perform an alignment between the “with-helmet” and “without-helmet” scans. The software produces dense point clouds of the external head surface, including the scalp and facial regions, as the full visible surface is captured during 3D scanning. In the next step the software converts the point clouds to a mesh model. Following scanning, the facial areas were removed during processing to preserve participant anonymity. The remaining scalp surface was then used for co-registration and subsequent analysis. Finlly to construct the head model, we used the fiducial landmarks to align the co-registered surface to the structural MRI.

**Figure 3.**
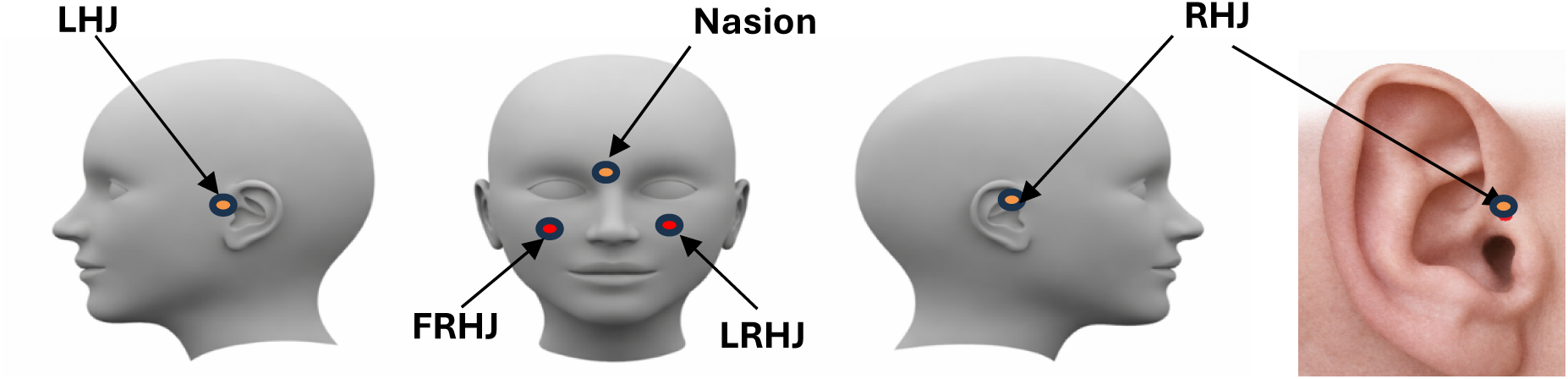
The fiducial landmarks. Position of the Nasion, Left Preauricular Point (LPA), Right Preauricular Point (RPA) along with two “fake fiducials” called Fake Right Preauricular Point (FRPA) and Fake Left Preauricular point (FLPA) are shown above. These points are marked on the participant’s face using a skin friendly marker.

For alignment to the participant’s structural MRI, the processed scalp surface mesh was co-registered to the MRI-derived scalp surface using the MNE python GUI. This co-registration was initialised using the fiducials (nasion and bilateral Helix-Tragus Junction points) identified on the MRI, followed by surface-based refinement using the full scalp geometry.

In the FieldLine OPM system, digitisation is performed with a Polhemus electromagnetic system using the Brainstorm software. A file is created to store the head shape and fiducial points. During the processing stage, OPM-FLUX uses this information to create a ‘device to head transformation’ file to be used during source modelling. The digitisation using the Polhemus is first carried out outside the MSR (without helmet). In addition to standard head shape points and anatomical fiducials (nasion [NA], left and right Helix-Tragus Junction points (LHJ and RHJ) Fig. 3), two additional reference points are recorded: FLHJ, and FRHJ because the Helix-Tragus Junction points (LHJ and RHJ) are partially or fully obstructed by the OPM helmet during scanning, they cannot be reliably digitised in the “with-helmet” condition. These FRHJ and FLHJ points are referred to as “fake” fiducials as they are not anatomical landmarks in the conventional sense. Instead, they are reproducible points selected on the skin surface that remain visible and accessible when the helmet is worn. They serve as practical substitutes for the true fiducial points during helmet-head alignment. The positions of the anatomical points are shown in the Fig 3. These points are marked on the participants face using a skin friendly marker. During digitisation, these two fake fiducials along with the NA are recorded twice to improve reliability. Subsequently, with the helmet on, reference points (marked on the Fieldline helmet) are recorded and the same fake fiducial points along with the NA point (previously marked on the skin) are recorded again.

Hence in the Fieldline system, an affine transform is first calculated using these fiducial points and helmet reference points to align the helmet (device) coordinate system with the head coordinate system. This transform is then applied to re-reference the helmet data, including sensor locations (originally defined in device space), into head coordinates. Once all elements are expressed in head space, a new .fif file (copied from the original) is generated containing updated sensor locations in head coordinates, digitised head shape points, and anatomical fiducials (nasion, LHJ, RHJ). This produces a dataset fully aligned in head coordinates for subsequent MRI co-registration and forward modelling.

### 3.2 Degaussing

For the Cerca system it is recommended to degauss the shielded room prior to any recording. Degaussing removes any residual magnetisation from walls of the magnetically shielded room (MSR). The degaussing coils are installed inside the walls of the MSR and can be operated through the acquisition PC. Degaussing was not performed in the FieldLine OPM system, as the fixed helmet array ensured a stable magnetic environment and hence provided no measurable performance gain.

### 3.3 Field nulling

Field nulling is a procedure used in the OPM system to reduce residual magnetic fields inside an MSR after the degaussing process. This is typically achieved using large-scale field-nulling coils installed within or on the wall of the MSR. For the Cerca system the field nulling coils are managed by the c-coil software (ref?). In contrast, FieldLine system does not rely on the external room-level nulling coils. Instead, each FieldLine OPM is operated in a closed-loop configuration using on-board sensor compensation coils, which actively null the magnetic field local to the vapour cell.

Because both OPM systems were operated in a closed-loop mode with on-sensor compensation, explicit room-level field nulling was not performed in either of the cases. The closed-loop feedback significantly increases the effective dynamic range of the sensors, allowing reliable operation even in the presence of residual background fields. Nevertheless, applying additional room-level field nulling may be advantageous in experimental conditions involving participant movement, where larger field variations can be encountered.

### 3.4 Empty Room Recording

Before starting the participant recordings, it is recommended to carry out an empty room recording for 2-3 minutes. This helps identify potential sources of artefacts, such as nearby fans or large electrical equipment. It also provides a chance to spot any faulty or untuned sensors, which can then be disabled during the actual recordings.

### 3.5 Structural Imaging

For each participant the T1 weighted MRI scans are typically recorded to support the source modelling. It is recommended to use an MP-RAGE or FLASH scan (see https://surfer.nmr.mgh.harvard.edu/ for details). Here, the MRI data were acquired using an MP-RAGE sequence (Brant-Zawadzki et al., 1992).

## 4. Design of the Pipeline

The data analysis pipeline is illustrated in Fig. 2, highlighting the key steps of OPM-FLUX, while the specific procedures and steps are outlined in the sections below. The corresponding scripts for each step can be accessed on the OPM-FLUX website (https://www.neuosc.com/flux/) linking to the relevant GitHub repository.

### 4.1 BIDS conversion

The pipeline begins by importing the raw OPM data, which were stored in fif format and split. To organise the OPM data and make it compatible with common analysis tools, the data are converted according to the MEG-BIDS format (Niso et al., 2018). Before conversion, the data were down-sampled to reduce file size and computational analysis load. The Cerca system having a sampling rate of 1500Hz the data were down sampled to 750Hz and the Fieldline system having a sampling rate of 5000Hz were resampled to 1000Hz.

To avoid aliasing problems the data were low-pass filtered at 250 Hz using MNE-Python before the down-sampling. This was achieved by applying a band-pass filter (0.1–250 Hz). To remove slow drifts, we used a linear-phase finite impulse response (FIR) filter with a Hamming window. The filter was applied in a zero-phase (forward–reverse, non-causal) configuration to prevent phase distortion. Filter length was 49,501 samples (33.0 s), yielding ∼53 dB stopband attenuation, with transition bandwidths of 0.10 Hz (lower; −6 dB at 0.05 Hz) and 62.5 Hz (upper; −6 dB at 281.25 Hz). During the BIDS conversion, essenfial metadata and file structure were preserved, while certain identifying information was anonymised to ensure compliance with the UK General Data Protection Regulation (UK GDPR). Data were anonymised using MNE default function, which clears subject-related metadata (info[’subject_info’]), removes experimenter identifiers, and resets measurement dates, without altering the MEG signal or acquisition parameters.

The BIDS format includes the trial related information which was recorded using the trigger lines in the Cerca/Quspin system. The trigger lines are identified and combined into a single digital trigger channel using a standard binary coding scheme. For the Fieldline system all the trigger information is stored inside a single stimulus channel called ‘di32’.

An event dictionary was created to map trigger values to their corresponding experimental conditions. These event markers are integrated in the BIDS structure. The event information is available inside the BIDS directory as a .tsv file. The snippet of the file is given in Fig. 5.

**Figure 4.**
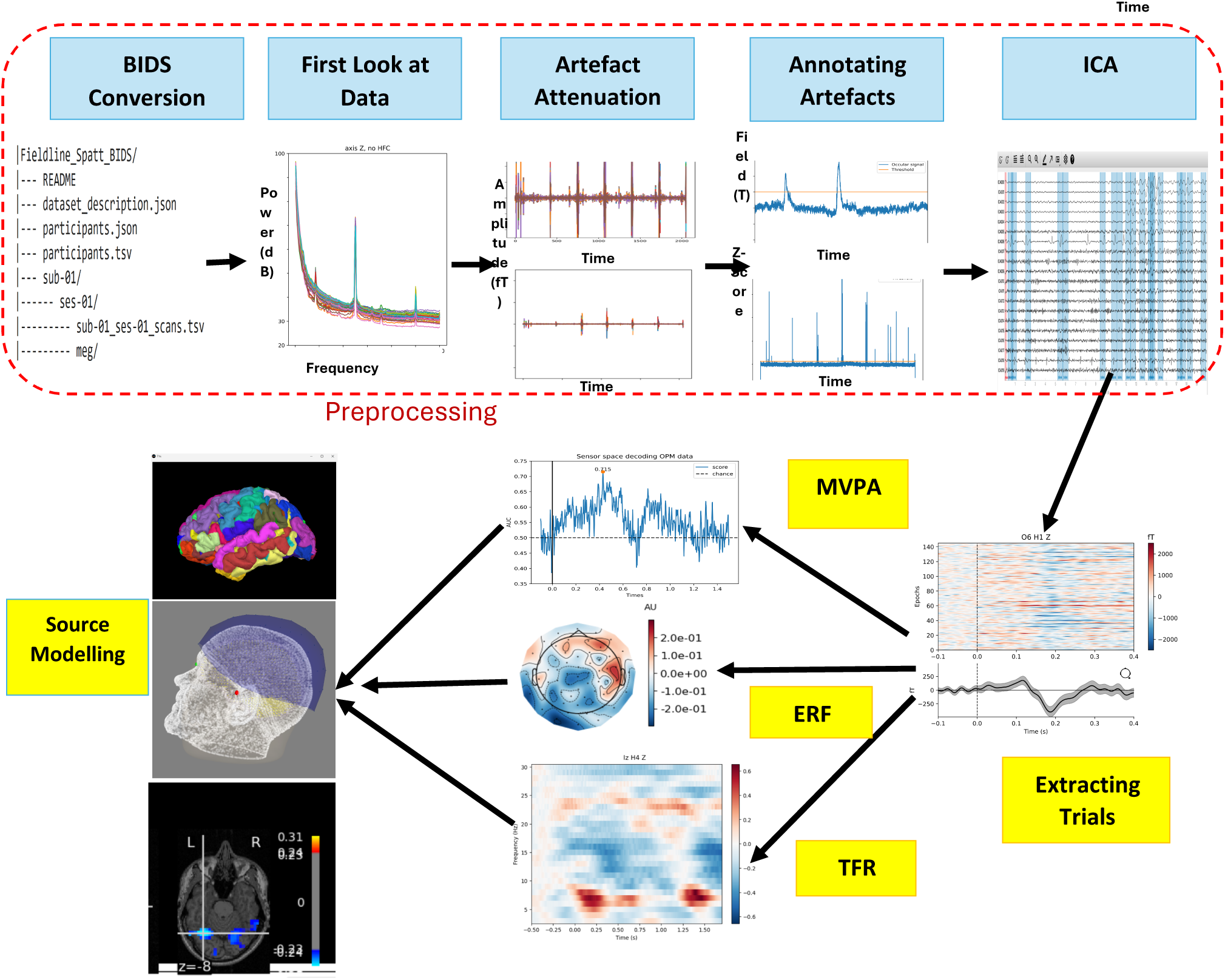
Overview of the OPM-FLUX processing pipeline. The workflow begins with raw OPM data undergoing down-sampling and BIDS conversion, followed by an initial inspection and artifact attenuation. Subsequent steps include artifact annotation and Independent Component Analysis (ICA) for advanced noise removal. After preprocessing, the cleaned data can be used for various analyses, including source modelling, event-related field (ERF) analysis, time-frequency representation (TFR), and multivariate pattern analysis (MVPA).

**Figure. 5.**
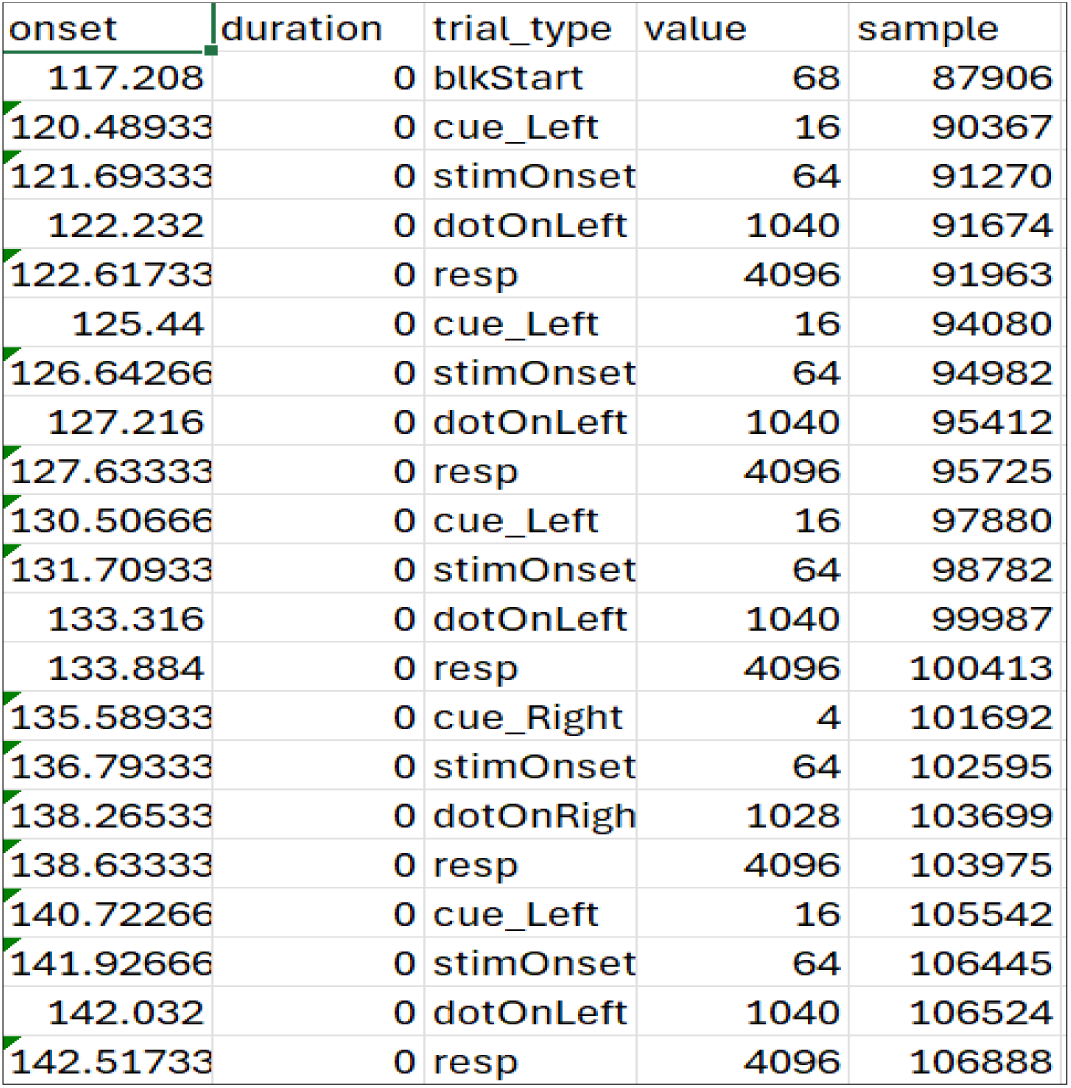
A snippet of the tab-separated values ( .tsv) file containing the event information after the BIDS conversion. As an example, value 16 reflects the trigger signifying *cue_Left .* The first occurrence is at sample 90367 time point 120.48933s. It is recommended to inspect the .tsv file to ensure the triggers are identified. Special care should be taken for triggers overlapping in time (e.g. a finger response trigger overlapping with a trial bases trigger).

### 4.2 Preprocessing

The raw OPM signal is contaminated with artefacts and noise arising from various sources like external magnetic field including power line interference, physiological sources like eye blinks, head movements, muscle artefacts etc, which must be attenuated. We here make these steps explicit.

#### 4.2.1 Removing faulty sensors

The preprocessing steps starts with visualisation of the data and the power spectra. In the Cerca system, OPM sensors record magnetic field data along three orthogonal axes, and therefore the power spectral densities (PSD) were estimated separately for each axis. In contrast, the Fieldline OPM sensors recorded magnetic field signals along a single axis only, being most sensitive to the radial field.

In the subsequent stage, the Power Spectra Densitometry (PSD) estimates are used to identify potentially faulty sensor. for this purpose, we plot the histogram of sensor PSDs and a threshold of PSD magnitude is selected from visual inspection. Sensors crossing a threshold are considered faulty and hence not included in the subsequent analysis. This is done separately for each of the three axes for the Cerca system.

The histogram plot of the power across all frequencies and all sensors, for each of the X, Y, and Z axes, is shown in Fig. 6 for the Cerca system. It can be observed that the power of nearly all sensors is largely confined within 39.5 dB. Therefore, this value was taken as a threshold, and any sensor with power exceeding 39.5 dB was discarded from the subsequent analysis.

**Figure. 6.**
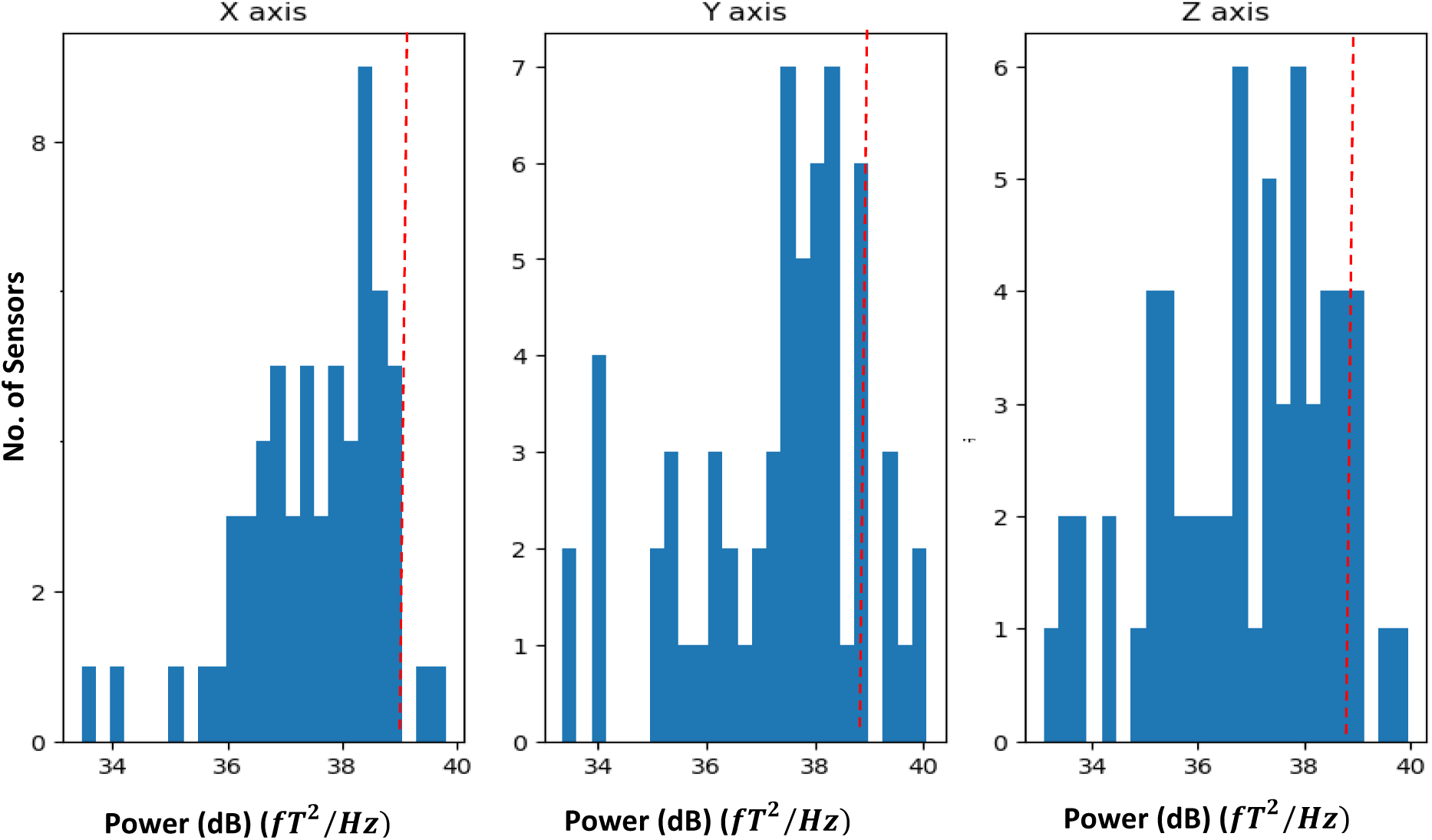
Histograms of the power averaged across frequencies plotted for the sensors for the X, Y and Z axes. Sensors with power *above 39.5 dB were considered faulty and discarded from further analysis*.

#### 4.2.2 Noise Reduction Algorithm (Homogeneous Field Correction)

Once the faulty sensors are removed, the raw data are subjected to homogeneous field correction (HFC). The HFC assumes the external interference as a homogeneous field and attempts to separate the signal from the interference. It estimates this homogeneous field by finding the best-fit constant field vector across the sensor array at each time point and projecting out the contribution from the sensor data. It is good to note that in the conventional MEG system some of the widely used noise removal techniques such as Signal Space Separation (SSS) (Taulu & Kajola, 2005); (Taulu & Simola, 2006) are applied to remove the noise in MEGIN systems. However those methods require precise measures of the sensor location and orientation and they apply a heavy regularisation which reduces the rank of the data considerably. Also, SSS is not optimal for multivariate approaches (Bezsudnova et al., 2024). Hence, we recommend using the homogeneous field correction (Seymour et al., 2022; Tierney et al., 2021) with aims at reducing interference from distant magnetic sources.

Fig. 7 shows and an example of the HFC analysis, demonstrating how the approach effectively attenuate noise in the data. The results are shown in both the time and frequency domains, highlighting the reduction in noise across each representation. Fig 7A and 7B show the raw data and the corresponding PSD respectively before applying the HFC. Fig. 7C and Fig. 7D shows the OPM signal applying the HFC. The mean power spectra before and after applying HFC is also shown in Fig. 7 E. This shows a ∼30% reduction of power reflecting the attenuation of artefacts.

**Figure 7.**
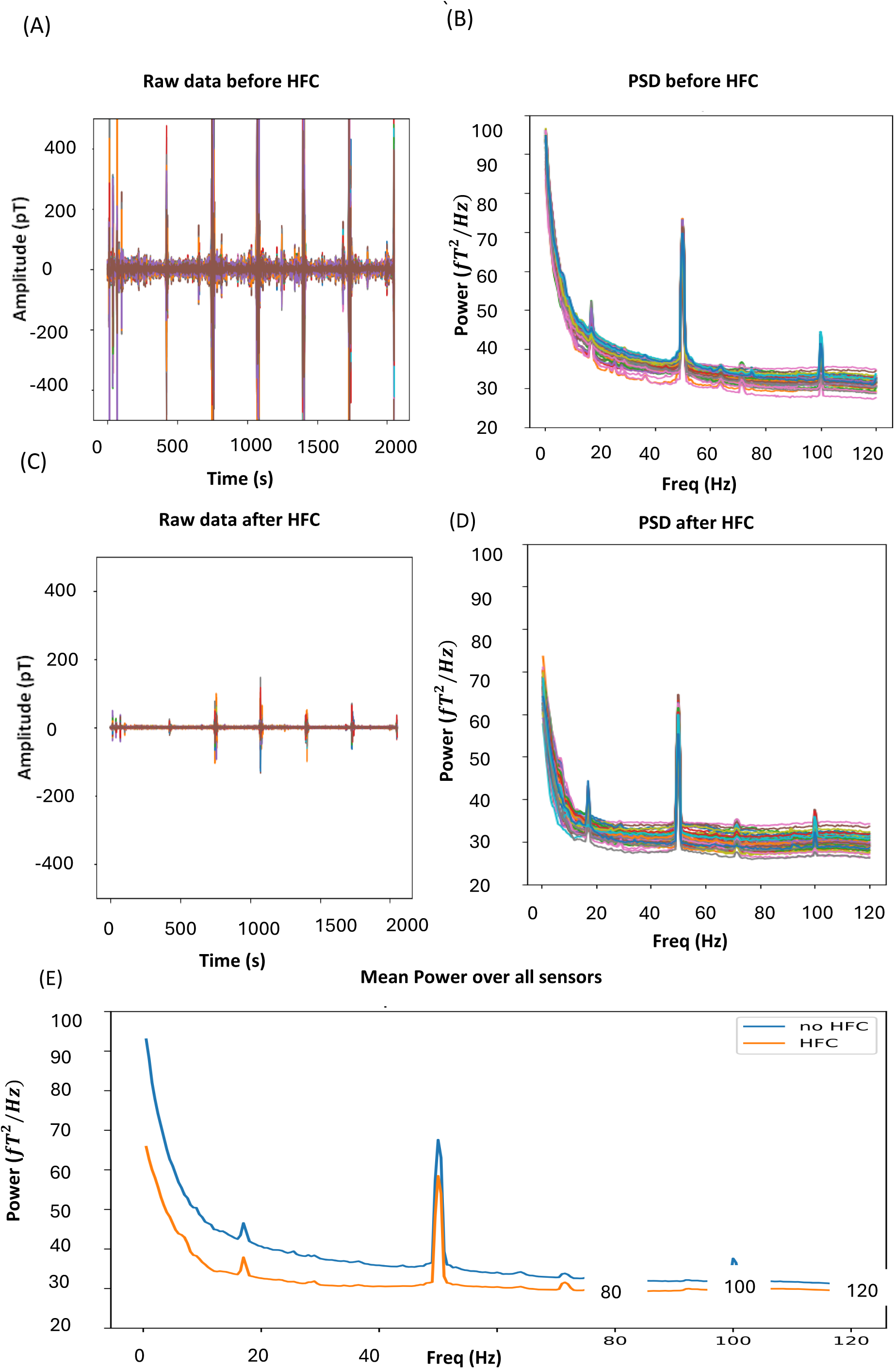
The figure illustrates the raw OPM signal and demonstrates how the HFC filter effectively reduces homogeneous magnetic noise. Images are given in both time and frequency domain. A) The raw signals before HF with the colours reflecting the different sensors. B) The power spectral density before the HFC. Note the 50 Hz line noise and an 18 Hz artefact most likely from the ventilations system. C) The raw signal after the HFC were applied. D) The power spectra after the HFC were applied. E) The mean power spectra over all sensors before (blue) and after applying HFC (orange). For clarity, only the radial components (Z-axis) of the tri-axial sensor are shown.

#### 4.2.3 Annotation of Artefacts

The next step is the identification and annotation of artefacts. Head and body movements, eye blinks and saccades, muscle activity, and malfunctioning sensors are major sources of artefacts in the OPM recordings. Movement artefacts are typically in the lower frequency ranges and strongly correlated across a large number of sensors, whereas malfunctioning sensors usually introduce abrupt jumps or discontinuities in the signal.

Eye blinks and saccades are identified primarily in the frontal OPM sensors, which are most sensitive to ocular activity. Ocular artefacts predominantly occur in the low-frequency range; therefore, the raw data are band-pass filtered between 1–10 Hz to enhance their visibility.

Automatic detection of ocular artefacts is performed using MNE-Python’s annotation functions, which flags time segments exceeding a predefined amplitude threshold. This threshold is selected based on visual inspection of the filtered data in the interactive MNE browser, ensuring that typical eye blink and saccade events are captured while minimizing false detections. The threshold may vary across participants. When available, eye-tracker data may also be used to corroborate the timing of blinks and saccades.

##### Annotation of Muscle Artefacts

Muscle artefacts are characterized by high-frequency broadband activity, typically in the range of 110 – 130 Hz. To detect these artefacts, the raw data are band-pass filtered at 110 – 130 Hz. Muscle artefact annotation is carried out using automated routines in MNE-Python based on the z-score of the filtered signal. The z-score threshold is determined empirically by visually inspecting the distribution of high-frequency activity across sensors and time. A threshold is chosen such that periods with abnormally elevated broadband power (indicative of muscle activity) are reliably identified while preserving physiological brain signals. A z-score value of 3 is chosen here.

It is generally advised to first mark any artifacts in the data and then decide whether to exclude the affected trials based on the research question. For example, muscle artifacts can interfere with measuring neuronal gamma-band activity because their frequency ranges overlap, but they may be less of a concern for slower event-related fields such as the N400m which are dominated by lower-frequency components.

#### 4.2.4 Independent Component Analysis

Independent Component Analysis (ICA) is a method used to separate a multivariate signal into statistically independent components (Hyvärinen & Oja, 2000; Lee, 1998; Makeig et al., n.d.) OPM-FLUX pipeline uses it to attenuate contributions from ocular and cardiac artefacts (Vigario et al., 2000). To reduce the computational overhead, OPM-FLUX recommends to bandpass signals within 3 – 30 Hz and then down-sample the data to 250Hz before applying ICA. The algorithm *fastICA* (Hyvärinen & Oja, 2000) is used to identify the independent components.

The ICA algorithm produces individual components on topographical maps along with their respective time series. The ocular and cardiac artefacts are identified by manually inspecting both topographies and time series. Typically, the artefacts are reflected in 3-5 components. We recommend not to reject more components unless they are associated with a known artefact. It is important to mention that cardiac artefacts might get attenuated due to the homogeneous field correction applied previously, and hence may not result on a component in the ICA analysis.

After identifying ICA components associated with artifacts, their influence can be attenuated by “projecting them out” of the data. It is important to apply these projections to the original raw data rather than the down-sampled version.

The choice between attenuating ocular artifacts with ICA or rejecting trials with blinks or saccades depends on the research question. For example, in studies of covert spatial attention, it is crucial that participants maintain central fixation; therefore, trials containing saccades may be excluded. In contrast, for studies on e.g. auditory perception, attenuating saccadic artifacts with ICA but not rejection may be more appropriate.

### 4.3 Extracting trials according to condition-specific events

The next step in the pipeline is to extract the relevant data segments according to the analysis requirements. Pre- and post-stimulus intervals are determined based on the research question. During trial extraction, previously annotated artefacts are applied to reject trials, while the event labels used for segmentation are defined during BIDS conversion based on trigger information.

### 4.4 Event-related fields

Event-related fields (ERFs) are brain responses that are time-locked with specific sensory, cognitive, or motor events. These responses reflect time-locked neural activity triggered by distinct stimuli. ERFs are widely used in cognitive and clinical neuroscience to study the timing of brain processes and to examine how the brain reacts to specific tasks or sensory inputs. When computing event-related fields (ERFs), it is advisable to be guided by the established guidelines for event-related potentials (ERPs) (Luck, 2014; Woodman, 2010) The trial length and filter settings should be determined based on the specific experimental design and research question.

In the dataset examined, the ERFs reflect the response to visual stimuli. We demonstrate how to illustrate the temporal patterns of the ERFs and generate topographical maps.

Here, the epochs are extracted for the condition with respect to the onset of the left and right cue onset. Prior to the computation of ERFs, the OPM data were band-pass filtered in the 0.1 – 30 Hz range to attenuate slow drifts and high-frequency noise and cropped to the −0.1 to 0.4 s interval from stimulus onset. The filtering was performed using a non-causal finite impulse response (FIR) filter, designed with a Hamming window and a filter length of 331 samples. This resulted in a 0.0194 passband ripple and 53 dB stopband attenuation. To avoid filter edge artefacts arising from signal discontinuities at epoch boundaries, it is essential to apply the filtering to the continuous data before epoch extraction. Following filtering, epochs were averaged across trials and baseline correction was applied by subtracting the mean signal within a 100 ms pre-stimulus interval. Since the QuSpin sensors are triaxial, ERF topographical maps are generated separately for each of the three independent orientations. A representative example of ERF is given in Figure 8, where the ERF (corresponding to the onset of the *Left Cue* visual stimuli) recorded from one OPM sensor (the radial z-direction) located over the right occipital area of the visual cortex is shown. The time evolving topographical plots corresponding to the same ERF is also shown. Note the dipolar pattern reflecting the visual evoked response over posterior sensors.

**Figure. 8.**
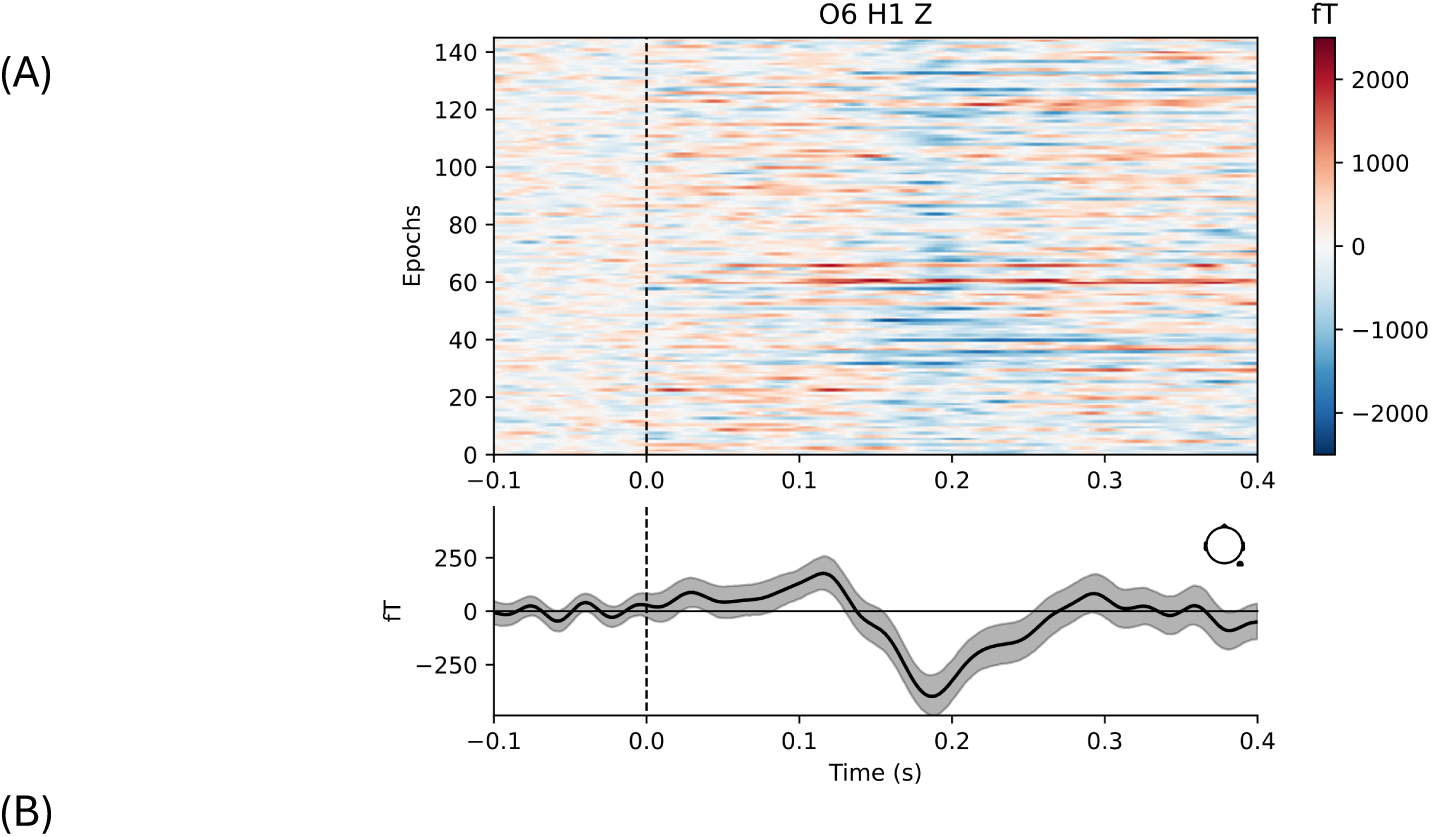
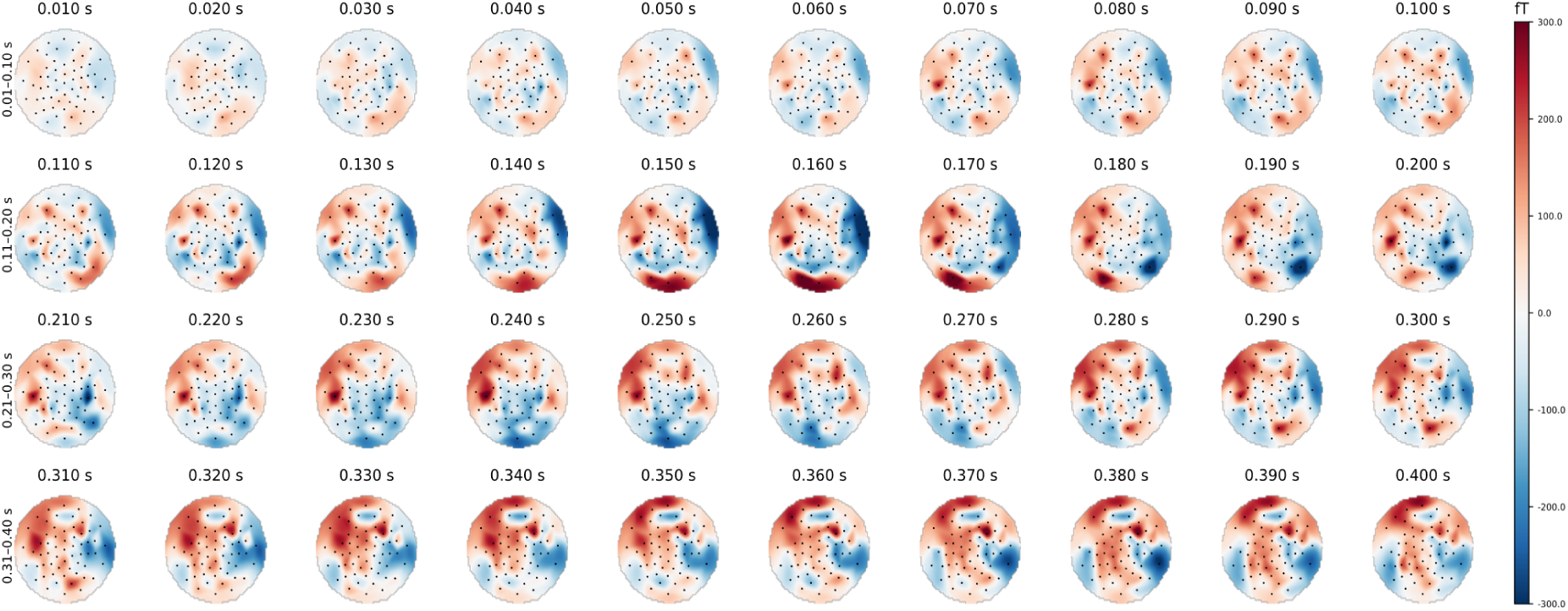
(A) A representative event-related field (ERF) corresponding to ‘Left Cue’ recorded from a sensor (O6 H1, z-axis) positioned over the visual cortex. (B) The topographical plot of the ERF over time. Note the posterior distribution reflecting a dipole in visual cortex.

### 4.5 Time-frequency representation of power

Neuronal oscillations have been implicated in supporting human cognition (Buzsáki, 2006; Jensen et al., 2007). Modulation of brain oscillations can be quantified in the time-frequency domain in trial-based paradigms using Morlet wavelet transformation, a sliding time-window Fourier Transform (Cohen, 2019) and bandpass filtering followed by a Hilbert transform.

These approaches produce comparable result when the parameters are adjusted appropriately (Bruns, 2004)In the current examples, we show how to use a sliding window Fourier transform approach to estimate the time-frequency representations of power.

From practical experience, it is known that low-frequency oscillations, such as those in the theta and alpha ranges tend to be narrow-band (*ΔF* ≈ 2 *Hz*), while gamma-band activity (60–90 Hz) is typically broader-band (*ΔF* ≈ 10 *Hz*). To account for these differences, we recommend using separate parameter settings when quantifying slow and fast brain oscillations.

For slow oscillations (<30 Hz), a sliding time window of *ΔT* = 500 ms is recommended. The window length is defined in cycles using *n_cycles* = freqs / 2, which then results in the window duration of *ΔT* = 500 ms across frequencies. Spectral estimation is performed using a DPSS (Slepian) tapers, where the time–bandwidth product controls both the number of tapers and the achievable frequency smoothing. Setting *time_bandwidth* = 2.0 corresponds to a time–bandwidth product of *2ΔTΔF* and results in a single DPSS taper being used (*N* = *time_bandwidth* − 1 = 1). For a window length of *ΔT* = 0.5 s, this configuration yields an effective spectral smoothing of approximately *ΔF* ≈ 3 Hz, which follows from the relation *ΔF* ≈ *3 / (2ΔT)* and reflects the minimum achievable frequency smoothing when one DPSS taper is applied. An example figure is given in Fig 9A, which shows the time-frequency representation of the alpha-band (10–12 Hz) power decrease after stimulus onset, with the corresponding topographical distribution.

**Figure 9.**
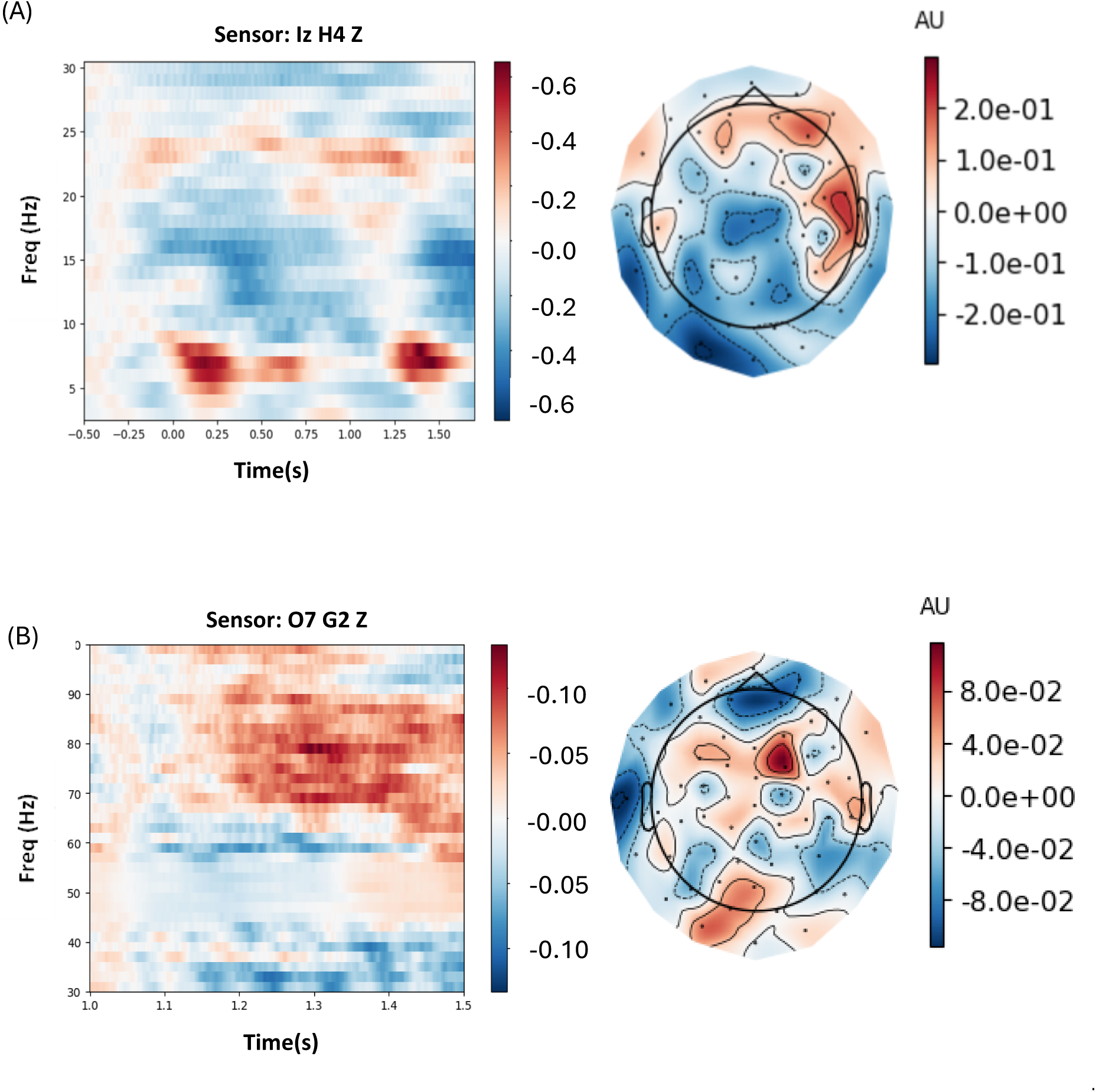
An example of the time–frequency representation of power for different frequencies and topographical maps, computed using the FLUX pipeline. (A) a time-frequency representation of power and corresponding topological map demonstrating a decrease after stimulus onset in the 10–12 Hz alpha band, and extending into the 20 Hz beta band. The early response at 0.2s in the 5 – 8 Hz theta reflets the spectral content of the event-related field. (B) A clear increase in gamma-band power (70–90 Hz) that can be observed following the onset of the gratings (t = 1.2 s). The power estimates were obtained using a multi-taper method to enhance spectral smoothing. A topological map corresponding the gamma-band power is also shown. Within the FLUX pipeline, specific recommendations are provided for optimising the analysis of oscillatory power modulations across both slow (<30 Hz) and fast (>30 Hz) frequency ranges.

For higher frequencies (>30 Hz), a full multitaper spectral estimation approach (Percival & Walden, 1993) is recommended, using a shorter sliding window of *ΔT* = 250 ms. In this case, multiple DPSS tapers are applied within each time window to improve the spectral estimation. The desired frequency smoothing (*ΔF*) is specified via the time–bandwidth product, and the corresponding number of tapers follows from this choice. With the chosen time-bandwidth, this results in 3 DPSS tapers being applied. According to the relation *N < 2ΔTΔF*, using *N* = 3 tapers corresponds to a minimum achievable spectral resolution of approximately *ΔF* ≈ 6 Hz. An example figure is shown in Fig.9B which shows the time-frequency representation showing gamma-band (70–90 Hz) power increase following grating onset (t = 1.2 s), with the corresponding topographical distribution.

Time-frequency of power can be assessed either by applying a baseline correction or by directly comparing experimental conditions. For a 500 ms window, a suitable baseline period is −500 to −250 ms, avoiding overlap with post-stimulus activity. However, the baseline interval may be adjusted depending on the experimental design, with the main principle being to exclude post-stimulus responses.

Modulations in spectral power can be quantified as a relative change from baseline, calculated as *P*_*rel*_ = (*P*_*stim*_ − *P*_*base*_)/*P*_*base*_(Pfurtscheller & Lopes da Silva, 1999). When comparing two conditions, A and B, this can be quantified as *P*_*rel*_ = (*P*_*A*_ − *P*_*B*_)/(*P*_*A*_ + *P*_*B*_) which provides a normalised measure of difference across participants. Alternative normalization methods, such as the log-ratio (*P*_*log*_ = *log*(*P*_*B*_ / *P*_*A*_)) or t-/z-score transformations, can also be applied but should be used cautiously since the power value distributions are positively skewed.

### 4.6 Multivariate pattern analysis

Multivariate pattern analysis (MVPA) is a widely used approach for analysing distributed brain signals across multiple sensors. Traditional univariate methods, although well established, typically rely on averaging across trials to improve signal-to-noise ratio. While this strategy is effective for detecting robust evoked responses, it may reduce sensitivity to subtle and temporally precise neural patterns that are essential for investigating dynamic cognitive processes such as the allocation if spatial attention. Trial averaging can obscure fine-grained temporal signatures that evolve rapidly over time. In contrast, MVPA operates on single-trial data and exploits spatially distributed patterns of activity across sensors or source space. This allows time-resolved decoding of neural representations and provides a powerful framework for characterising rapidly changing brain dynamics. The core principle of MVPA is to identify condition-specific representational patterns rather than focusing on amplitude changes at individual sensors. For OPM-MEG, which can involve many sensors with multiple measurement axes, MVPA is particularly well suited. By leveraging the rich multichannel information inherent in OPM recordings, MVPA offers strong potential for capturing distributed neural representations that may not be detectable using conventional univariate approaches.

Two types of MVPA are often used: classification-based decoding and representational similarity analysis (RSA). In classification-based decoding (Stokes et al., 2015; van de Nieuwenhuijzen et al., 2013), activity patterns across sensors are used to distinguish between experimental conditions. At each time point, sensor values are combined into a feature vector, and a classifier such as linear discriminant analysis (LDA) or a support vector machine (SVM) is trained to differentiate between brain states. The trained model is then tested on independent data to evaluate how accurately neural patterns predict the condition. This approach can be applied to binary or multi-class problems.

In contrast, RSA quantifies the similarity between neural response patterns across conditions. These neural similarity structures are then compared to model-based similarity matrices to assess how well different theoretical representations explain the observed brain activity (Cichy et al., 2014; Cichy & Oliva, 2020)

A key consideration in MVPA is the selection of an appropriate classification method. Given the wide range of machine learning algorithms available, it is important to choose a method that is well suited to the structure of neuroimaging data and the objectives of the experiment. In the OPM-FLUX pipeline, we adopt a classification approach to distinguish between two conditions specifically, brain states associated with attending to the ‘left versus the right cue’ based on distributed patterns of brain activity. For this purpose, we use a Support Vector Machine (Cortes & Vapnik, 1995) with a radial basis function (RBF) kernel (Wainer & Fonseca, 2021). This approach is widely used in MEG research due to its robustness, strong generalization performance, suitability for high-dimensional data, and its ability to perform well with a limited number of training samples while maintaining relatively low computational complexity. Implemented within the MNE-Python framework, it provides a reliable and efficient method for decoding distributed brain activity patterns.

In this pipeline, the data were first bandpass filtered from 1 to 45 Hz and down sampled to 125 Hz to reduce computational load. This was done before segmenting the continuous signal into shorter trials. A time window from −0.1 to 1.5 s around cue onset was selected for analysis. For classification, tools from the Scikit-Learn library (Pedregosa et al., n.d.) were used. The data were organized as a 3D matrix (trials × sensors × time) and transformed into a 2D format suitable for machine learning. The features were standardized to have zero mean and unit variance. A processing pipeline was created to handle the transformation and classification steps. Time-resolved classification was performed using a support vector machine, applied independently at each time point. A 10-fold cross-validation approach was used, where the classifier was trained on 90% of the trials and tested on the remaining 10%, repeated across 10 iterations. Performance was measured using the Area Under the Curve (AUC), and results were plotted over time. An example of MVPA on OPM data using the OPM-FLUX pipeline is shown in Fig. 10, where left versus right attention trials were classified with a SVM classifier. The highest classification accuracy was observed around 400 ms after target onset, with an AUC of approximately 0.72.

**Figure. 10.**
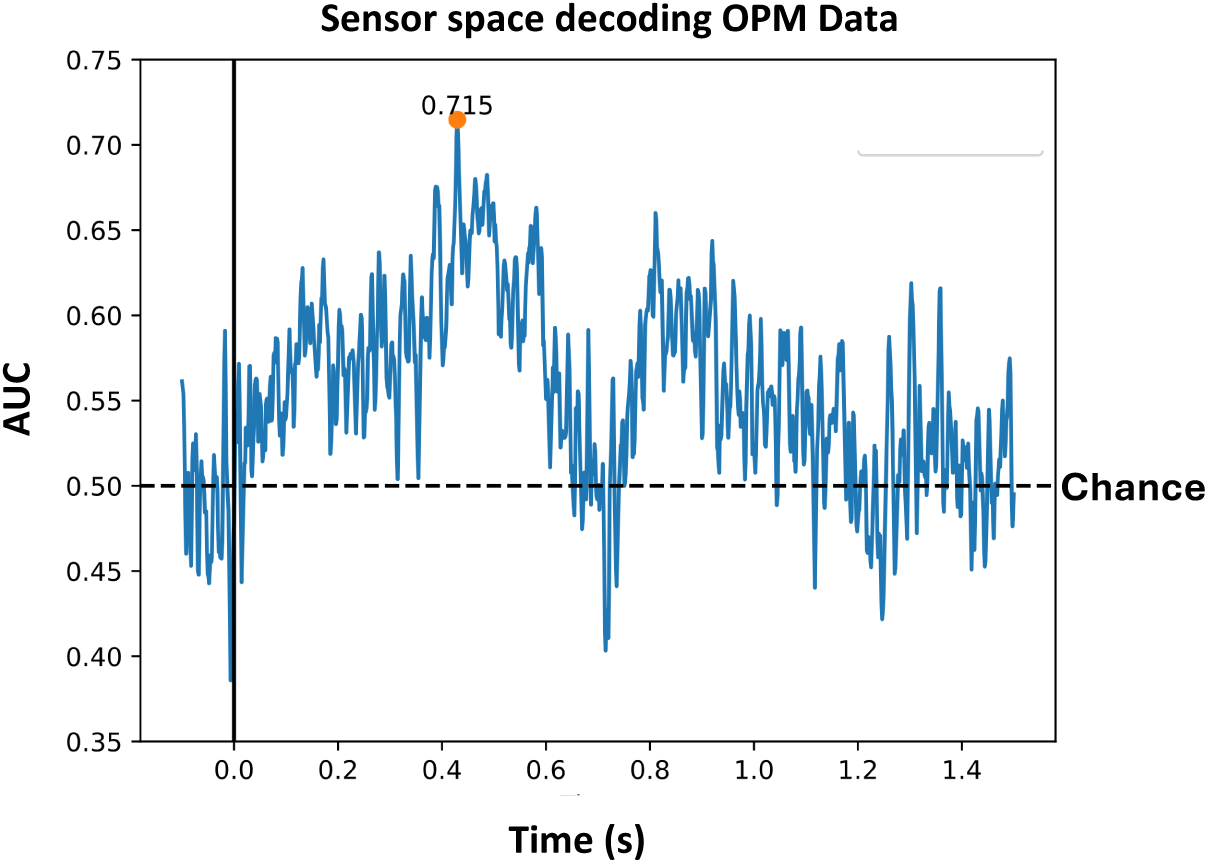
An example of multivariate pattern analysis (MVPA) performed on OPM data using the OPM-FLUX pipeline is shown above. In this illustration, trials corresponding to left versus right attentional allocation were classified using a support vector machine (SVM) with a radial basis function (RBF) kernel.

### 4.7 Source modelling

Source modelling involves estimating where brain signals are generated and can be done using different approaches. In the past, dipole modelling was used to estimate the location of a few discrete sources, but nowadays distributed source methods are standard. Here we focus on localising oscillatory brain activity using a beamforming approach based on Dynamic Imaging of Coherent Sources(DICS) (Gross et al., 2001). The source modelling is performed using the following major steps in sequence.

#### Step1: Acquire and preprocess the structural MRI

High-resolution T1-weighted MRI is acquired for each participant and converted to Neuroimaging Informatics Technology Initiative (NIfTI) format. The T1 image is processed in FreeSurfer to produce cortical and skull/scalp surfaces. These surfaces form the anatomical basis for a realistic head model that will be used in the subsequent forward-model computations.

#### Step2: Construct scalp/brain surfaces and BEM

The FreeSurfer reconstruction is used to generate the brain surface. This surface forms the basis of the head model used for forward modelling. A single-shell Boundary Element Model (BEM) is then created. This model defines the geometry of the head needed to calculate how brain activity produces the magnetic signals detected by the sensors. The resulting BEM is subsequently used to compute the forward solution required for accurate source reconstruction.

#### Step3: Co-registration with anatomical landmarks

The sensor array-to-head transformation, based on digitised scalp landmarks obtained from the 3D scan as well as landmarks on the sensor array, is performed using the inbuilt Cerca software as discussed in section3.1. The next step is to align the head (in sensor space) with the MRI anatomical space based on the anatomical landmarks (nasion, left and right Helix-Tragus Junction points). This spatial alignment is used for computing the forward solution. To perform this alignment, the MNE-Python coregistration GUI is used, which allows us to visually match the digitized head shape and fiducial points to the MRI surface.

#### Step4: Define the volumetric source space (grid)

A volumetric source space is created by tiling the brain volume with a regular three-dimensional grid. For OPM-MEG we use a grid spacing of 5 mm.

#### Step5: Compute the lead-field matrix (forward solution)

Using the BEM, the sensor array-to-head transformation, the aligned source grid, and the measured sensor positions, the lead-field matrix (forward solution) is computed. The lead-field matrix describes how activity at each grid points of the lead field project to the MEG sensors. This forward solution is a key component of the source reconstruction and is reused across all subsequent inverse-method analyses.

#### Step6: Source modelling

Dynamic Imaging of Coherent Sources (DICS) is used to localise changes in oscillatory brain activity within specific frequency bands. DICS is a frequency-domain beamforming method that estimates source-level power by combining the forward (lead-field) model with cross-spectral density (CSD) matrices computed for predefined time windows and frequency ranges. In the present analysis, CSDs are estimated for a pre-cue (−0.7s to −0.2 s) and a post-cue interval (0.1 to 0.6 s), using the same window length and taper settings as in the TFR analysis of the FLUX scripts.

The CSD matrix captures frequency-specific covariance between all sensor pairs and therefore preserves both amplitude and phase relationships at the frequency band of interest. In DICS, the CSD serves the same role as the covariance matrix in time-domain beamformers (Van Veen et al., 1997) but in the frequency domain. CSD matrices can be estimated using a multitaper approach for the selected frequency band and time window.

To enable valid comparisons across conditions, a common spatial filter is constructed by averaging the CSD matrices from all conditions together. This combined CSD provides the covariance structure used to compute a single set of spatial filters, ensuring that differences in source-level power reflect genuine neural changes rather than condition-specific filter differences. DICS calculates a volumetric source grid covering the whole brain.

Beamforming methods such as LCMV and DICS can exhibit depth-related bias due to differences in sensor sensitivity and noise characteristics across the brain (Gross et al., 2001; Van Veen et al., 1997). As a result, superficial sources may exhibit higher estimated power than deeper sources, independent of true neural activity. Importantly, since the present analysis focuses on comparing conditions across different time intervals within the same participants, rather than on absolute source power estimates or comparisons across special locations. Because the same forward model, source space, and spatial filters are applied consistently across all conditions any residual depth-related bias affects both conditions similarly and therefore does not systematically influence the contrasts of interest.

In conventional SQUID-MEG systems, an explicit noise covariance matrix derived from empty-room recordings may be used for spatial whitening and improving the sources estimates. However, in OPM-MEG estimates of the noise covariance matrix is complicated by the fact that the sensor array is moving in the MSR. As a result, empty-room recordings do not accurately reflect the noise characteristics present during the experimental data acquisition. For this reason, no separate noise covariance matrix was applied here. Instead, regularisation of the cross-spectral density (CSD) matrix and careful estimation of the effective data rank were used to ensure stable and well-conditioned spatial filter construction. This approach follows established practice in OPM-MEG beamforming analyses, where the use of empty-room covariance is often infeasible or inappropriate.

The spatial filters are computed using the forward model and the combined CSD. To stabilise the inversion, Tikhonov regularisation is applied by adding a small constant λ (5% of the sensor power) to the diagonal of the CSD matrix. This diagonal loading increases the smallest eigenvalues, ensuring that the matrix is well-conditioned and can be reliably inverted. It also prevents near-zero eigenvalues from dominating the inversion and amplifying noise, resulting in more stable and robust spatial filter estimates.

Because covariance or CSD matrices can be rank-deficient, particularly in OPM-MEG due to preprocessing steps such as homogeneous field correction or ICA, the effective data rank is estimated prior to inversion. A truncated pseudo-inverse is then computed using singular value decomposition (SVD). For a data matrix X with rank k, the decomposition is given by

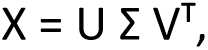

and the truncated pseudo-inverse by

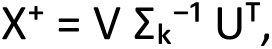

where only the first k singular values of Σ and the corresponding singular vectors in U and V are retained. This approach ensures stable and accurate computation of the spatial filters.

For each source location (grid point), the source orientation is optimized to maximize output power by selecting the orientation associated with the largest eigenvalue of the local source power matrix. The resulting spatial filters are then applied to condition-specific CSDs to obtain source-level power estimates, which are mapped onto the individual structural MRIs for anatomical interpretation.

An example of alpha-band (8 – 12 Hz) and gamma band (70 – 90Hz) source modelling within the cue target interval 0.1s to 0.6s window and 1.2 – 1.7s window in which the gratings occur respectively is shown in Fig. 11.

**Figure.11.**
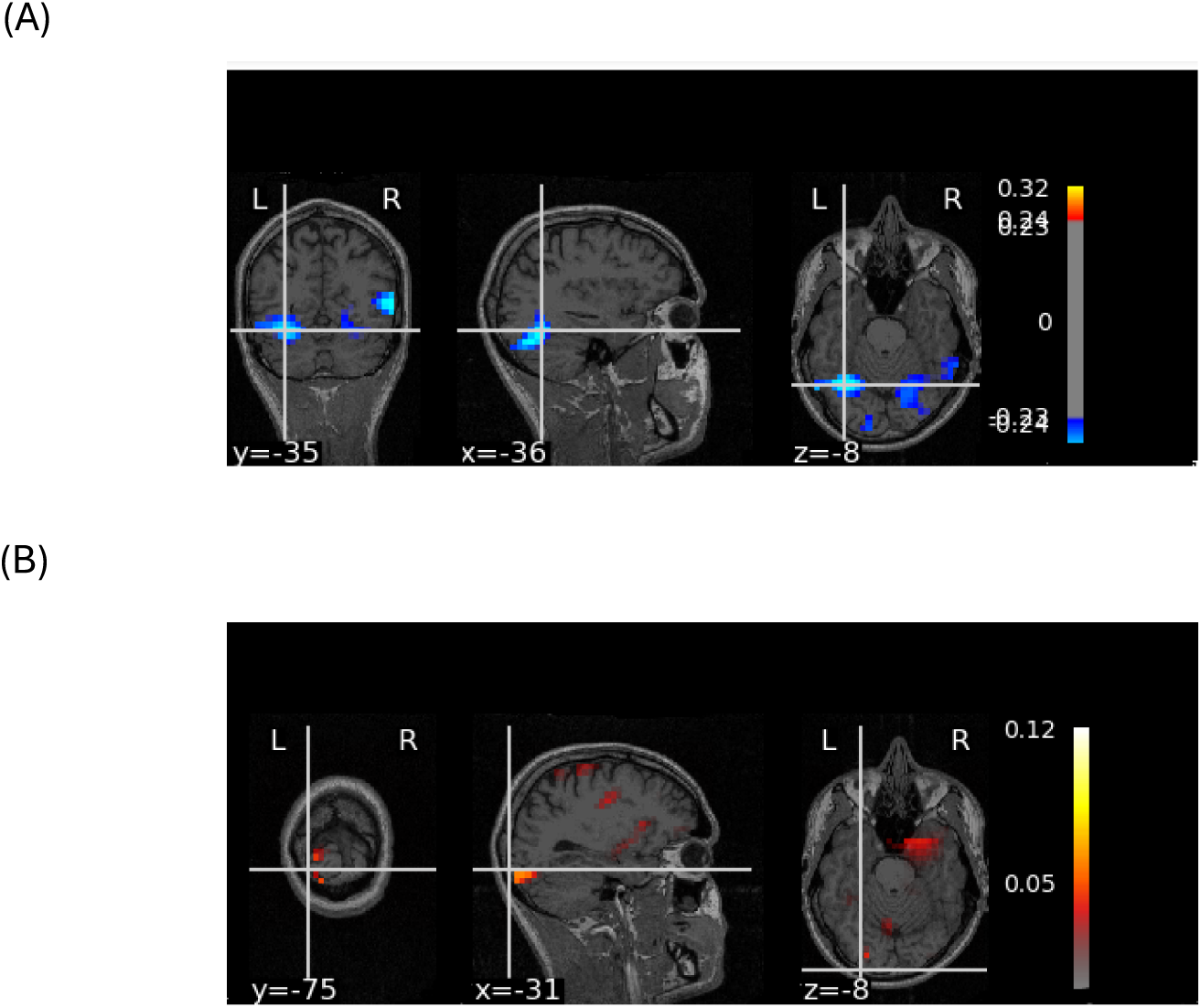
(A) An example of source modelling for alpha-band activity (8 – 12 Hz) within 0.1s to 0.6s time window B) The gamma band (70 – 90Hz) activity within 1.2 –1.7s. The analysis was performed using the DICS beamformer implemented in MNE-Python.

## 5 Discussion

The OPM-FLUX pipeline provides a dedicated framework for the analysis of MEG data acquired with optically pumped magnetometers (OPMs), with the primary goal of streamlining analysis procedures and promoting consistency in parameter selection. By offering a standardised workflow, OPM-FLUX enhances the reproducibility and transparency of OPM-based research and includes structured recommendations on which methodological details should be reported in publications and pre-registered protocols, supporting best practices in open science.

Another key feature of the pipeline is the inclusion of detailed tutorials accompanying each script and processing step. These tutorials explain the technical background of the methods used, clarify their relevance for OPM signal processing, and guide users through parameter choices, enabling informed adaptation of the analysis to specific datasets. As a result, users can follow the complete workflow, from raw recordings to final outputs, while gaining a clear understanding of each stage.

Developed using openly available test datasets from the Cerca and FieldLine systems, OPM-FLUX also serves as a teaching and training resource suitable for self-guided learning, workshops, and formal educational settings. In this way, the pipeline not only supports rigorous and reproducible data analysis but also lowers the entry barrier for researchers new to OPM-MEG.

### 5.1 Novelty of the toolbox

As OPM technology is relatively recent, only a limited number of dedicated toolboxes and processing pipelines are currently available for OPM-based cognitive neuroscience research. Many of the existing solutions are MATLAB-based, including OPM-specific processing tools as well as established packages such as SPM(Litvak et al., 2011). More general-purpose toolboxes like FieldTrip and Brainstorm have also been extended to support OPM data, providing flexible frameworks for MEG analysis. In addition, MNE-Python offers several OPM-specific processing methods, however, a fully integrated end-to-end pipeline still needs to be put together by the user; we here offer a solution

The OPM-FLUX is based on Python-based programming, which is one of the most widely accepted languages for data science and machine learning. The OPM-FLUX pipeline makes use of MNE-Python, an open-source Python toolbox for neuroscience studies, which provides great flexibility for cognitive neuroscience research and supports many state-of-the-art approaches such as multivariate pattern analysis and classifier design, both of which are essential components of the proposed pipeline. Beyond these features, OPM-FLUX takes advantage of Python’s large collection of scientific and machine-learning libraries, including NumPy, SciPy, and scikit-learn, making it easy to connect with state-of-the-art computational tools and to extend the pipeline as needed. The pipeline works smoothly across different operating systems and does not require expensive software licenses, making it accessible to a wider research community. Because both Python and MNE-Python are open source, OPM-FLUX supports reproducible and transparent research and benefits from continuous improvements driven by the global user community. In addition, its modular and flexible design allows users to customize the workflow for different types of studies, such as experimental or clinical research, and to easily adapt to future advances in technology and data-analysis methods.

### 5.2 Future of OPM-FLUX

As the OPM technology continues to evolve rapidly, with advancements in quantum sensors (ranging from alkali to helium-based types) and new approaches such as camera-based head tracking and reference sensors for motion regression, the OPM-FLUX pipeline will evolve to keep pace with these developments. It will be regularly updated to integrate the latest methods and technologies in the field. At the same time, we will ensure that the core pipeline remains simple and stable, while more advanced or experimental developments are implemented as modular extensions that can branch out from the core workflow.

In the near future, one goal is to integrate an eye-tracking system alongside OPM recordings. This integration will open exciting new research possibilities for studying the relationship between brain oscillations and eye saccades. Another immediate objective is to incorporate head movement data directly into the processing pipeline to further reduce motion-related artifacts and improve data quality. In addition, future versions of OPM-FLUX will include group-level statistical analyses at both the sensor and source levels, using non-parametric cluster-based permutation approaches. All corresponding codes and analysis modules will be integrated into the OPM-Flux scripts, ensuring a unified and accessible framework for data acquisition and processing.

Looking ahead, OPM-FLUX also aims to support batch processing capabilities by integrating with the OHBA Software Library (OSL-ephys) framework (van Es et al., 2025). This will make large-scale data analysis more efficient and user-friendly, ensuring that the pipeline remains a flexible, scalable, and future-ready tool for the growing OPM research community.

## 6 Conclusion

This paper advocates the use of the open-source standardized pipeline OPM-FLUX, developed specifically for cognitive neuroscience studies using OPM-MEG. Since the OPM technology is still new, it is essential to standardise the general processing steps associated with OPM experiments. Having a dedicated end-to-end pipeline tailored for OPM ensures a coherent and efficient workflow that addresses the unique characteristics of OPM data.

The OPM-FLUX pipeline is built entirely on open-source resources, promoting reproducibility, transparency, and accessibility in research. Its primary goal is to guide users in defining a consistent analysis path for their projects amid the wide range of options and providing a clear structure and standardised approach to encourages well-documented, transparent publications.

Moreover, the pipeline supports pre-registration practices by offering concrete recommendations on what aspects of the analysis should be reported. Beyond research applications, OPM-FLUX can also be effectively used in educational and training settings, helping to raise the methodological standards in the field of cognitive neuroscience and fostering a new generation of researchers familiar with best practices in OPM-MEG analysis.

## Acknowledgements

This work was supported by Wellcome Trust Discovery Award (grant number 227420) and the NIHR Oxford Health Biomedical Research Centre (NIHR203316).

The views expressed are those of the author(s) and not necessarily those of the NIHR or the Department of Health and Social Care. The Oxford University Centre for Integrative Neuroimaging was supported by core funding from the Wellcome Trust (203139/Z/16/Z and 203139/A/16/Z).

We also acknowledge Dr. Robert Seymour for his support and assistance with the OPM data collection at the Oxford Centre for Human Brain Activity (OHBA), University of Oxford.

## Data Availability Statement

Codes and datasets that are freely accessible via the Neural Oscillation Group website (https://www.neuosc.com/flux).

